# Mondrian Abstraction and Language Model Embeddings for Differential Pathway Analysis

**DOI:** 10.1101/2024.04.11.589093

**Authors:** Fuad Al Abir, Jake Y. Chen

## Abstract

In this study, we introduce the Mondrian Map, an innovative visualization tool inspired by Piet Mondrian’s abstract art, to address the complexities inherent in visualizing biological networks. By converting intricate biological data into a structured and intuitive format, the Mondrian Map enables clear and meaningful representations of biological pathways, facilitating a deeper understanding of molecular dynamics. Each pathway is represented by a square whose size corresponds to fold change, with color indicating the direction of regulation (up or down) and statistical significance. The spatial arrangement of pathways is derived from language model embeddings, preserving neighborhood relationships and enabling the identification of clusters of related pathways. Additionally, colored lines highlight potential crosstalk between pathways, with distinctions between short- and long-range functional interactions. In a case study of glioblastoma multiforme (GBM), the Mondrian Map effectively revealed distinct pathway patterns across patient profiles at different stages of disease progression. These insights demonstrate the tool’s potential to enhance downstream bioinformatics analysis by providing a more comprehensive and visually accessible overview of pathway interactions, offering new avenues for therapeutic exploration and personalized medicine.

## I. Introduction

In functional genomics and proteomics, effective visualization and analysis of complex biological data are crucial for elucidating the intricate networks of gene expression, protein interactions, and pathway activities [1], [2]. Traditional visualization tools, however, often struggle with these complexities, leading to challenges in clarity and comprehensiveness [3].

To surmount these challenges, we introduce the Mondrian Map, a novel visualization tool that integrates multiple dimensions of pathway analysis—pathway enrichment, crosstalk, and clustering—into a single, coherent framework. This tool not only facilitates a deeper understanding of biological networks but also supports a broad range of downstream analyses critical for advancing both research and therapeutic development.

Drawing visual inspiration from Piet Mondrian’s abstract art, the Mondrian Map uses fixed colors and grid structures as metaphors to enhance visual clarity and structure. However, the tool’s scientific rigor is driven entirely by the underlying biological data. Data-driven algorithms determine the layout, connections, and sizes of the squares within the map, ensuring that the visualization is both aesthetically appealing and biologically meaningful.

Traditional tools like Cytoscape [4], VisANT [5], STRING [6], and Graphia [7] have significantly advanced the study of biological networks but often fall short in handling complex, large-scale networks. For instance, Cytoscape’s node-edge representations can become cluttered with increasing network size [8], while GeneTerrain faces challenges related to noise sensitivity and lacks mechanisms for visualizing pathway crosstalk or clustering [9]. These limitations can obscure the functional relationships within biological networks, impeding the extraction of actionable insights.

The Mondrian Map addresses these limitations by offering clear representations of pathway dynamics and interactions. Its innovative visualization framework uses size and color coding to reflect pathway activity and significance, while colored lines indicate pathway crosstalk, differentiating between functional interactions to reveal potential pathway synergies or antagonisms. Language model embeddings inform the spatial arrangement of pathways, preserving neighborhood relationships and facilitating the identification of functionally related pathway clusters. This clustering is especially crucial in complex diseases like glioblastoma multiforme (GBM), where understanding pathway interplay is essential for identifying therapeutic targets.

Beyond mere network visualization, the Mondrian Map’s utility extends to functional genomics and proteomics. It enables the identification of key regulatory pathways and their interactions, offering deeper insights into protein function, modifications, and interactions within the context of pathway enrichment and crosstalk.

Previous work in systems-scale analysis and drug repositioning has highlighted the need for advanced visualization tools capable of integrating comprehensive pathway insights [10]–[13]. Furthermore, the critical role of pathway analysis in systems biology underscores the indispensability of such tools in unraveling complex molecular interactions [14]. Additionally, in the realm of personalized medicine, understanding individual variability in disease progression and treatment response is enhanced by the ability to visualize and interpret complex biological networks [15]. The Mondrian Map leverages its innovative framework to provide a comprehensive tool for researchers, facilitating the integration of pathway analysis into personalized treatment strategies.

In summary, the Mondrian Map represents a significant advancement in bioinformatics, functional genomics, and systems biology. By amalgamating pathway enrichment, crosstalk, and clustering into an intuitive visualization, it provides a powerful tool for analyzing and interpreting complex biological data, with the potential to significantly impact downstream bioinformatics analyses and advance personalized medicine.

## II. Methods

Using rectangular shapes, primary colors in bold blocks, and a strict grid structure, Mondrian is remembered for achieving a universal and harmonious visual language with his abstraction framework [16]. One of his original artwork is presented in supplementary file, appendix A. Harnessing these spatial features possesses an enormous potential for providing an intuitive representation of intricate scientific data. While this approach enhances comprehension, its implementation demands multiple stages of data processing particularly for complex biological networks. In this section, we described the detailed development process of our visualization tool, how we leveraged language modeling to generate pathway embeddings and subsequently address the essential data processing steps required for creating the visualizations.

### A. Language Model Embeddings for Pathways

#### 1) Selection of Language Models

We utilized a diverse set of language models to generate pathway embeddings which captures intricate neighborhood and interaction relationships. The models utilized include all-MiniLM-L6-v2 (MiniLM) [17] and all-mpnet-base-v2 (mpnet) [18], whose embeddings were produced using the Sentence Transformer library [19]. Additionally, LLM2Vec-Mistral-7B-Instruct-v2mntp (Mistral-7B) [20] and LLM2Vec-Meta-Llama-3-8BInstruct-mntp (Llama-3-8B) [21] were used, utilizing the llm2vec framework, which transforms large, decoder-only language models into bidirectional encoders. This transformation involves enabling bidirectional attention, masked next token prediction, and unsupervised contrastive learning, effectively creating rich, context-aware text embeddings [22].

In contrast, the Sentence Transformers (a.k.a. SBERT) adapts the pre-trained BERT architecture into siamese and triplet network structures to derive semantically meaningful embeddings efficiently, greatly reducing computational demands while preserving high accuracy in semantic similarity tasks [19]. The specific attributes of these models, such as the number of parameters, vocabulary size, and embedding dimensions are detailed in the Table I to contextualize their use in our analysis.

#### 2) Prompt Generation Techniques

We have generated the prompts to create the pathway embeddings in two phases. In the first phase, we used the Meta-Llama-3.1-8B-Instruct model to summarize the descriptions of the pathways, enhancing clarity and precision. Initially, the model removes alphanumeric identifiers, URLs, and external references to focus on the essence of the content. Then, for extensive descriptions, it distills the text to emphasize key functions and the pathway’s biological significance. From brief or one-liner descriptions, we synthesized concise summaries grounded in recent scientific literature, avoiding speculative content. The output encapsulates the summarized pathway description in a coherent paragraph not exceeding 300 words.

In the second phase, we generate structured prompts across four distinct categories, each targeting a specific aspect of pathway data, designed to be fed into language models listed in Table I for subsequent embedding generation.

**Type 1. Gene Symbol**. Lists up to 100 gene symbols sorted by their significance in the pathway.

**Type 2. Gene Description**. Augments gene symbols from Type 1 with it’s descriptions.

**Type 3. Pathway Name**. Simplifies the prompt using the pathway name only.

**Type 4. Pathway Description Summary**. Utilizes concise summaries generated in the first phase.

These prompts are crafted to guide the language models in generating high-quality embeddings that accurately reflect the complex dynamics of genetic and pathway interactions. Details on the prompts for embedding generation, including specific instructions added for llm2vec and inference parameters, are documented in the supplementary file, appendix B.

#### 3) Efficacy Evaluation of Embeddings

We employed the pathway embeddings along with their two-dimensional projections (UMAP and t-SNE) in a multi-class classification setting to evaluate the effectiveness of the embeddings. A standard Support Vector Machine (SVM) from Scikit-learn library [23] was applied without any hyper-parameter tuning to assess the inherent data patterns and structural properties.

Pathway annotations were sourced from WikiPathways [24], typically characterized by low-level details. We consolidated these detailed annotations into broader categories—such as classic metabolic pathways, disease pathways, signaling pathways, and regulatory pathways—using ontology data from BioPortal [25]. In instances where pathways corresponded to multiple categories, we assigned a single label randomly to mitigate overlaps.

### B. Mondrian Map Generation Algorithm

Following the characteristics and elements of Mondrian abstraction, we present a novel algorithm designed to generate the visualizations using scientific data. The algorithm necessitates several preprocessing steps of the raw data to utilize entity attributes and relationships (see Fig. 1 (left)). This can be structured into two tables which are presented in Fig. 1 (center) with pseudo data. The first table encompasses entity attributes such as entity IDs, fold changes, adjusted pvalues, and the layout co-ordinates whereas the second table should have entity relationships such as pair of entity IDs that shares a relationship. With these data tables, we can create the Mondrian Maps shown in Fig. 1 (right), where entities with different states and relationships are distinctively color-coded.

**Fig. 1.**
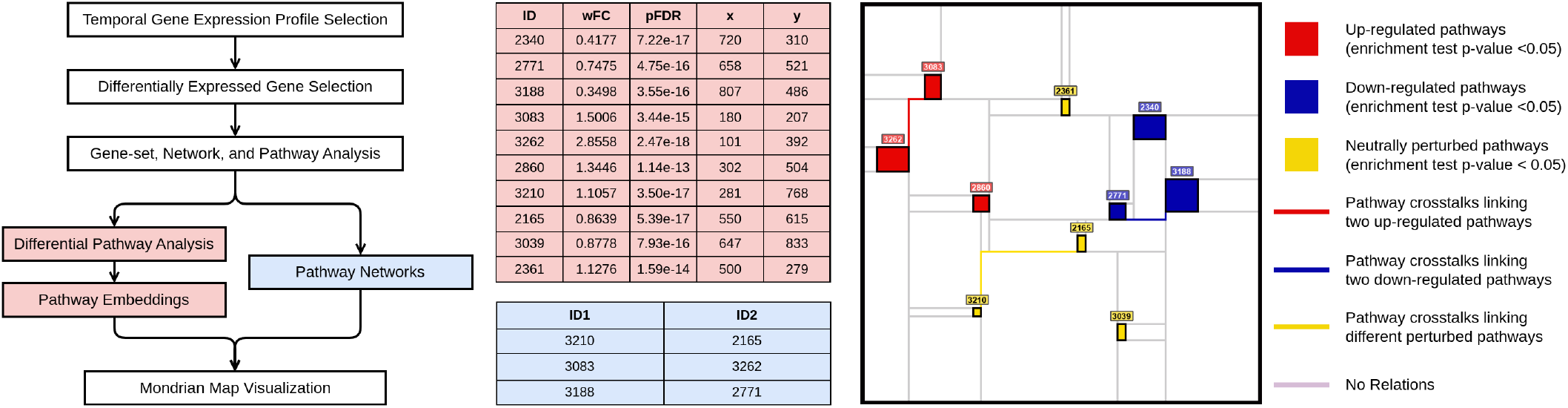
Data preparation flow chart for Mondrian Map (left). Getting the high-level insight from the Mondrian Map (right) is very intuitive compared to the tabular form of data (center). To facilitate that, effective color scheme for the entities and relationships are essential (right).

Essential attributes such as area, color, and central point coordinates are required to represent each entity as a rectangle whereas, the relationships between the entities are illustrated using lines that link the corners of the corresponding rectangles. The definition and significance of these attributes are subject to variability contingent upon the specific use case and the underlying scientific inquiry. In our case study, area of the rectangles are proportional to fold change of the pathways and the color depends upon fold change and adjusted p-values. For intuitive understanding of the Mondrian Map generation algorithm, we have depicted the key steps in Figure 2. The procedure to create a Mondrian Map is described as follows:

**Fig. 2.**
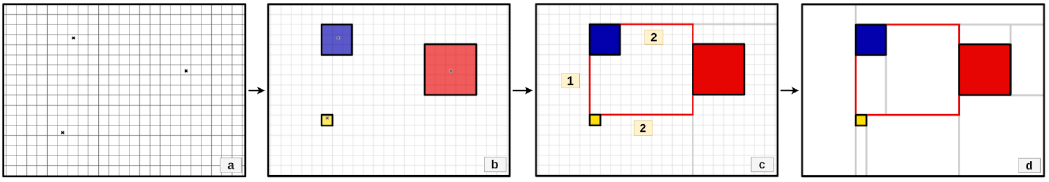
Implementing Mondrian Map involves 4 key steps: (a) Initiating a grid-based canvas with specified point coordinates, (b) generating rectangles to symbolize entities, (c) illustrating red lines to establish relationships between the entities, and (d) populating the canvas with additional gray lines.

**Step 1. Creating Canvas**. Initialize an empty canvas featuring a grid system with the central points (x, y) of all entities, depicted in Fig. 2(a).

**Step 2. Visualizing Entity Attributes**. Utilizing the central points and the designated area, rectangles are generated by approximating the number of rows and columns of grid-blocks surrounding the points and colored according to other attributes (Fig. 2(b)). The approximation of the point and area is deliberately imprecise to maintain adherence to the grid system.

**Step 3. Visualizing Entity Relationships**. Relationships between entities are established by identifying the two closest vacant corners of their rectangles and drawing red straight lines (Type 1) for vertically or horizontally aligned entities, or L-shaped lines (Type 2) for others (Fig. 2(c)).

**Step 4. Mimicking the Art Style**. The last step focuses on the art-style, drawing vertical and horizontal gray lines from L-shaped lines (Fig. 2(c)) and each vacant rectangle corners and retaining only the shorter line to enhance visual clarity depicted in Fig. 2(d). These lines are for illustration only and hold no significance in the analysis.

#### C. Data Preparation for Case Study

We employed longitudinal gene expression Glioma dataset to demonstrate the effectiveness of Mondrian Map generation. Filtering differentially expressed genesets from gene expression profiles of distinctive patient pools, we conducted geneset, network, and pathway analysis to obtain pathway regulatory networks. Subsequently, we generated utilized language model embeddings to find distinctive patterns in Mondrian Maps. This section details the essential data preparation steps required for Mondrian Map generation. We have also noted down the entity counts for each steps along with example tables in appendix C for transparency and reproducibility.

#### 1) Dataset Introduction

In our study, we utilized the Glioma Longitudinal AnalySiS (GLASS) dataset [26]. The consortium aims to enhance understanding of glioma tumor evolution and identify potential treatment targets. The dataset covers a wide range of patients, from those with highly aggressive gliomas to milder cases. Most importantly, it is the only dataset, to our knowledge, that includes gene expression data both before and after interventions across the trajectory of glioma progression. With these unique characteristics, it perfectly aligns with the requirements for effective Mondrian Maps for our case study.

#### 2) Profile Selection Process

To analyze different levels of glioma aggressiveness, we selected three distinct patient profiles from the GLASS dataset: aggressive, non-aggressive, and a moderate baseline. The selection involved:

#### Initial Screening

We identified patients with at least three tumor samples (Tumor Primary (TP), and the first (R1) and second (R2) recurrences). Only 10 out of 201 patients met this criterion.

#### Aggressiveness Profiling and Patient Selection

From these 10, two were chosen based on clinical data to represent the most and least aggressive cases. Patient ID 0279, diagnosed with Grade IV Glioblastoma at 34, survived 11 months after multiple surgeries, representing an aggressive profile. Patient 0027, on the other hand, diagnosed with Grade II Astrocytoma at 30 and later developing Grade IV Glioblastoma in 80 months, representing a non-aggressive profile.

#### Baseline Cohort Establishment

We created a baseline profile by averaging (median) the gene expression values of four patients with moderate aggressiveness, who survived between 37 to 42 months. These profiles will facilitate a detailed comparative analysis using Mondrian Maps to identify significant patterns.

#### 3) Differentially Expressed Gene Selection

We refined our dataset by applying a threshold of 0.001 to exclude genes with nearly zero expression levels across all nine profiles, reducing the dataset to 11,090 genes for further analysis. We calculated Fold Change (FC) values for R1 and R2 relative to TP, denoted as R1 vs TP (or, R1/TP) and R2 vs TP (or, R2/TP). We then set thresholds for up-regulation (*>*= 1.5) and down-regulation (*<*= 0.5) to identify six sets of differentially expressed genes (DEGs). For the aggressive patient profile at the first recurrence (R1), we found 745 up-regulated and 923 down-regulated genes; at the second recurrence (R2), there were 985 up-regulated and 1101 down-regulated genes. The non-aggressive and baseline profiles also showed variable numbers of DEGs, listed in Table 1 in appendix C.

**TABLE I.** Comparison of Different Language Models.

| Model | # Params. | Vocab. Size | Embed. Dim. |
| --- | --- | --- | --- |
| MiniLM [17] | 22M | 30,522 | 384 |
| mpnet [18] | 109M | 30,527 | 768 |
| Mistral-7B [20] | 7B | 32,000 | 4096 |
| Llama-3-8B [21] | 8B | 128,256 | 4096 |

#### 4) Gene-set, Network, and Pathway Analysis

Utilizing six differentially expressed genes (DEGs), we performed a comprehensive gene-set, network, and pathway analysis (GNPA). This unravels the interconnected functions of genes and their implications for specific conditions or diseases. We accessed the Pathway, Annotated-list, and Gene-signature (PAG) Electronic Repository (PAGER) API to obtain comprehensive data on gene groups characterized by shared pathways, biological processes, and expression patterns under specific conditions or common downstream targets [27], [28].

We exclusively sourced data from WikiPathways using the API, considering only statistically significant PAGs (*p − value <* 0.05). As WikiPathways only stores pathway information, we have used PAGs and pathways interchangeably in this paper. Subsequently, we retrieved the m-type PAG-to-PAG co-membership network using the PAG IDs. The m-type relationship networks connect pairs of pathways with a significant number of shared genes [27]. A summary of retrieved pathway and pathway relationship network is presented in appendix C (Table 2).

#### 5) Differential Pathway Analysis

We have retrieved all the genes and their RP-scores associated in those pathways to calculate weighted Fold Change (wFC) using PAGER API. The RP-score, stands for rank prioritization-score, is used to rank genes within the pathway and calculated using Eq. 1 [27].

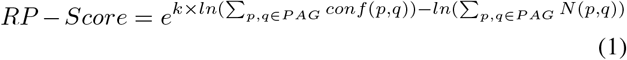

where p and q as gene indexes from the PAG, with a constant *k*(= 2), the term *conf* (*p, q*) represents the interaction confidence score ([0 *−* 1]) assigned by HAPPI-2 [29]. Additionally, *N* (*p, q*) takes on a value of 1 if gene *p* interacts with *q*, or 0, otherwise.

With the gene-level Fold Change (FC) values calculated previously and considering the RP-scores as weights (W), we have calculated weighted Fold Change (wFC) for each pathway using Eq. 2.

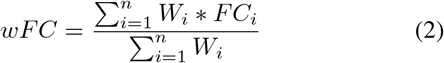

where *n* is the number of genes in the pathway, *W*_*i*_ is the RP-score of *Gene*_*i*_ and *FC*_*i*_ is the fold change value of *Gene*_*i*_.

#### 6) Pathway Representation

The visualization attributes in our study are adapted to the specific use case. We represent each pathway with a rectangle, where the size proportionate the absolute log2 of the weighted fold change (wFC), and position it on the canvas using t-SNE projections of language model embeddings. Pathways with *wFC* ≥ 1.25 are considered up-regulated and depicted in red, while those with *wFC* ≤ 0.75 are down-regulated and shown in blue. Pathways that do not meet these criteria are marked in yellow, unless they are deemed insignificant based on the adjusted p-value (*pFDR* ≥ 0.05), in which case they are depicted in black. Relationships are indicated with lines connecting rectangle corners: red for connections between up-regulated pathways, blue for down-regulated, and yellow for others; gray lines indicating no specific relationships (see Fig. 1 (right)). We also display the last four digits of the PAG ID at the top of each rectangle for easy reference and validation. At this stage, we have retrieved and processed all the data necessary for our visualization. The preprocessing steps are concisely summarized in the flow diagram shown in Figure 1 (left).

### III. Results

After thoroughly describing the prompting techniques to generate effective embeddings, the implementation of Mondrian Map and outlining the data processing steps in section II, this section dives deeper into a comprehensive analysis of the generated Mondrian Map visualizations resulting in the Glioblastoma case study. We highlight pathway regulations, relationships, distinctions and similarities among various profiles at different time points with 4 regions of interests: (a-d) from Fig. 4.

### A. Embedding Evaluation

We experimented with four language models using four different prompts to generate embeddings. Our analysis indicates that the t-SNE projection of Llama-3-8B embeddings, derived from the pathway description summaries, provides the most accurate projection. While Llama-3-8B embeddings based on pathway names achieved the highest classification accuracy at 73%, slightly surpassing the 72% accuracy of the description summaries, the latter’s richer data content led us to use its 2D coordinates for pathway localization. Figure 3 compares the 2D t-SNE projections across all methods, highlighting distinctive clusters, particularly in the Mistral-7B and Llama3-8B embeddings.

**Fig. 3.**
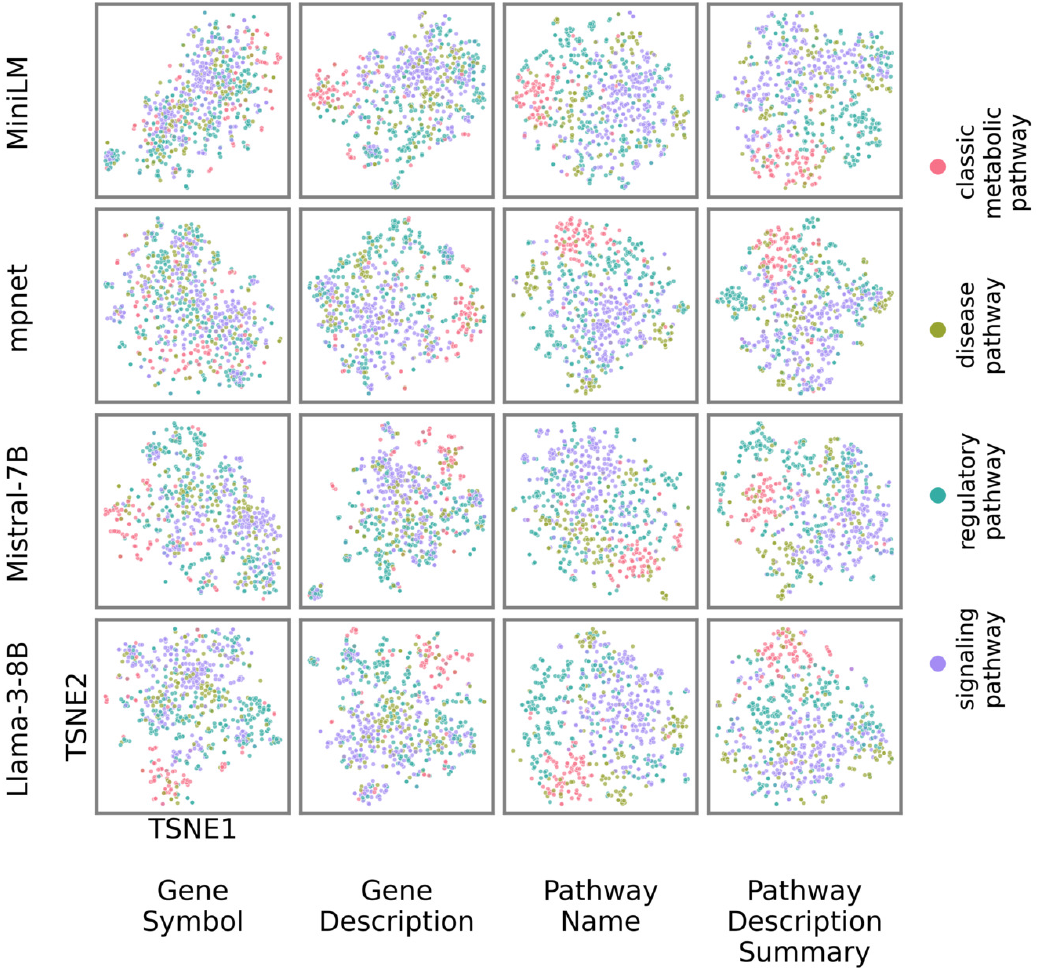
Comparison of t-SNE projection for the language model embeddings with different prompting strategies.

### B. Mondrian Map Analysis

Figure 4 illustrates the Mondrian Map visualizations of biological pathway activities across three glioblastoma (GBM) patient profiles, categorized by survival timelines: short-survival individual (*P*_*−*1_), baseline cohort (*P*_0_), and long-survival individual (*P*_+1_). These visualizations focus on the top 10 most significant pathway activity groups (PAGs) based on adjusted values. Each panel compares pathway regulation between initial recurrence (R1 vs. TP) and second recurrence (R2 vs. TP) for each cohort, highlighting four distinct regions (a-d) of interest. Below, we analyze the similarities and differences in pathway regulation between R1 and R2, emphasizing their roles in GBM progression and patient prognosis.

**Fig. 4.**
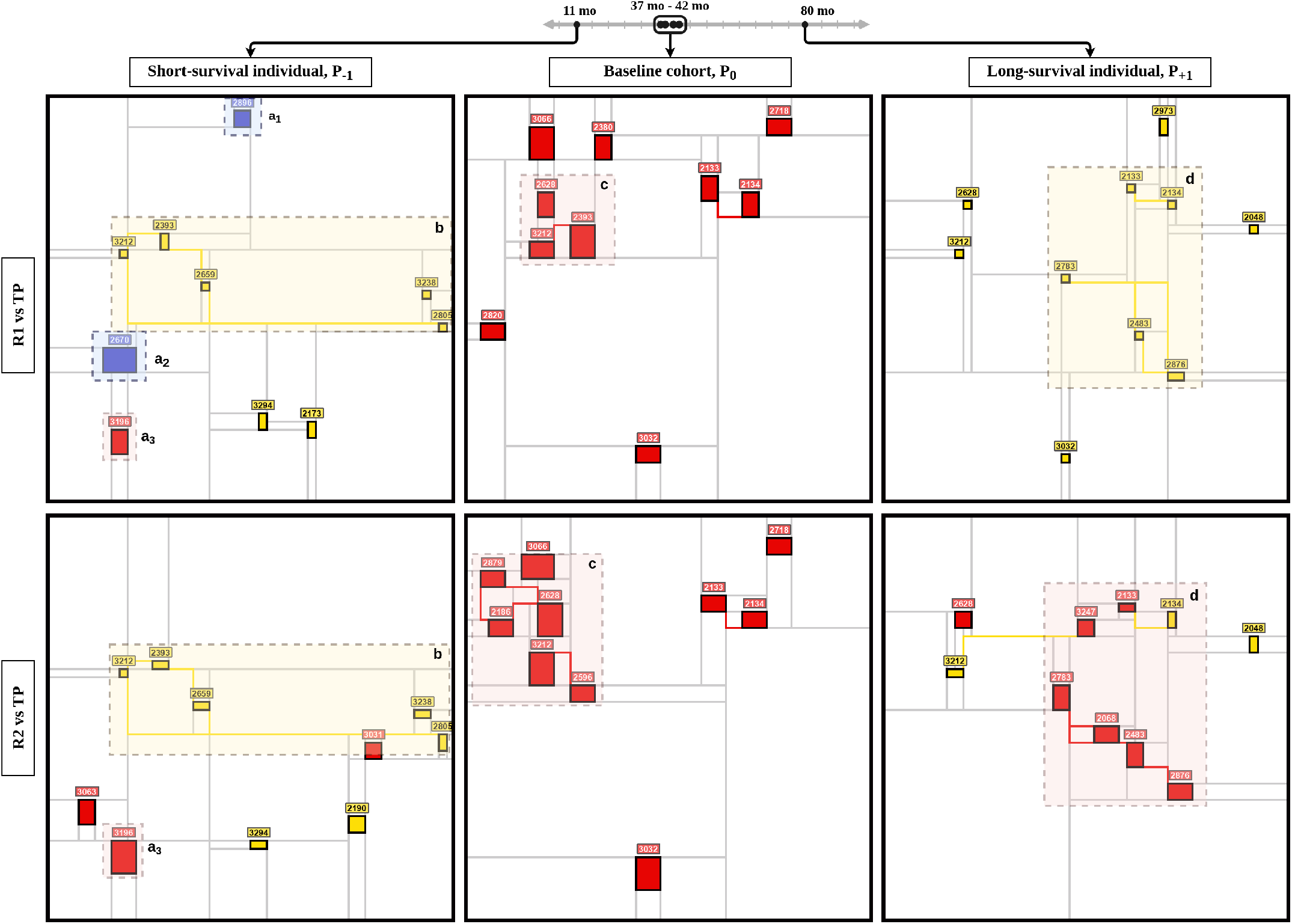
Comparative analysis of pathway dynamics across Glioblastoma patients using Mondrian Map. Profiles have been sorted in columns by the timeline of survival of the patients annotated at the top whereas the rows have the recurrences: R1 vs TP and R2 vs TP, respectively. Four regions of interest (a-d) are marked for further explanation

#### 1) Region a

In Region a, we observe both shared and distinct pathway activities between R1 and R2. The downregulation of G alpha (i) signaling (2896, sub-region *a*_1_) and TYROBP network in microglia (2670, sub-region *a*_2_) is evident in R1. This down-regulation suggests a sustained impact on oncogenic pathways and immune modulation [30]. However, the burn wound healing pathway (3196, sub-region *a*_3_) shows persistent up-regulation from R1 to R2. This sustained up-regulation indicates a continuous inflammatory and pro-tumorigenic environment, which may contribute to the aggressive progression observed in short-survival patients [31]. The down-regulation of G alpha (i) signaling in initial recurrence might lead to the unchecked activity of oncogenic pathways like ERK/MAPK and PI3K/Akt, which are critical for tumor survival and therapy resistance [30]. Similarly, the TYROBP network down-regulation suggests ongoing immune evasion mechanisms in the tumor microenvironment. The upregulation of the burn wound healing pathway reinforces the idea that glioma progression involves chronic inflammatory processes that are not easily reversed even after multiple recurrences [32].

#### 2) Region b

The spatial separation of pathways in region “b” within the embedding space, despite their significant crosstalk, suggests a complex interplay among them. This distribution highlights their distinct roles in cellular processes while simultaneously revealing their functional overlap in cancer biology, particularly in signaling pathways related to cell survival, proliferation, and adhesion. This indicates a tightly integrated network underlying cancer progression mechanisms.

The PI3K/Akt/mTOR signaling pathway (2805) plays a crucial role in glioblastoma (GBM) progression and therapeutic resistance. This pathway is frequently activated in GBM, contributing to tumor growth, survival, and invasion [33], [34]. Mutations or loss of the tumor suppressor PTEN, which normally inhibits this pathway, are common in GBM and lead to increased PI3K/Akt/mTOR signaling [35]. The pathway regulates various cellular processes, including cell cycle progression, metabolism, and protein synthesis through its downstream effectors like 4E-BP and S6K [34]. Inhibition of PI3K/Akt/mTOR signaling has shown promise in preclinical studies, with drugs targeting different nodes of the pathway demonstrating anti-tumor effects [36], [37]. Focal Adhesion Kinase (FAK, 2393) is a key mediator between focal adhesions and intracellular signaling. FAK activation promotes glioma cell invasion and migration through various signaling pathways, including PI3K-Akt [38] and FAK inhibitors have shown potential in treating various cancers, including glioblastoma [39]. While Malignant Pleural Mesothelioma (MPM, 3212) is a distinct cancer type from glioma/glioblastoma, it shares some similar molecular mechanisms. The PI3K-Akt-mTOR pathway is also activated in MPM, contributing to its aggressive nature which is evident by the crosstalks [40].

#### 3) Region c

Region c shows notable changes in pathway activities between R1 and R2, particularly in the baseline cohort. In R1, pathways such as focal adhesion signaling (2393) are prominently up-regulated, which aligns with the early aggressive features of GBM, including enhanced cell proliferation and migration [41]. However, by R2, there is a shift towards the up-regulation of pathways like EGF/EGFR (2879) and VEGFA/VEGFR2 (2628), highlighting the tumor’s adaptation towards mechanisms that support sustained growth and angiogenesis.

This shift from R1 to R2 indicates that as GBM progresses, there is a transition from pathways that drive initial tumor spread to those that sustain tumor expansion and survival under therapeutic pressure. The up-regulation of the EGF/EGFR pathway in R2, particularly involving the EGFRvIII mutant, suggests a growing reliance on growth factor signaling for continued tumor proliferation and therapy resistance [42], [43]. Similarly, the increased activation of the VEGFA/VEGFR2 pathway underscores the tumor’s need for enhanced vascular support as it expands [44].

#### 4) Region d

In Region d, significant changes between R1 and R2 are observed in the long-survival glioblastoma patient. Initially, in R1, the pathways related to ciliary function (2783, 2483, 2876) and miR-targeted genes in lymphocytes (2133) are relatively dormant. However, by R2, these pathways show substantial up-regulation, suggesting a shift towards enhanced cellular signaling and environmental sensing mechanisms that could facilitate tumor adaptation and evasion of immune responses [45].

The up-regulation of these pathways in R2 highlights the tumor’s evolving strategy to survive under increasingly hostile conditions, such as those imposed by treatments. The ciliary function pathways are particularly noteworthy, as recent studies have linked disruptions in primary cilia to GBM development and resistance to therapy [45]–[47]. The activation of miR-targeted genes in lymphocytes also suggests an ongoing interaction with the immune system, potentially contributing to immune evasion and sustained tumor growth.

#### 5) Summary of R1 vs. R2 Differences and Commonalities

The analysis of these regions across the Mondrian Map reveals that while some pathways are consistently dysregulated across both recurrences (e.g., PI3K/Akt/mTOR, FAK), others show significant shifts between R1 and R2 (e.g., miR-targeted genes in lymphocytes, ciliary function pathways). These shifts likely reflect the tumor’s adaptation to both its microenvironment and therapeutic pressures, highlighting potential windows for therapeutic intervention that could exploit these dynamic changes in pathway regulation.

## IV. Discussion and Conclusions

The Mondrian Map offers a transformative approach to visualizing complex biological networks, surpassing traditional methods such as heatmaps or node-edge representations. Unlike heatmaps, which focus on the regulation status of individual pathways without context, or traditional network visualizations that can become cluttered with excessive edges and nodes, the Mondrian Map excels in providing a holistic view of pathway interactions.

One of the key advantages of the Mondrian Map is its ability to visualize clusters of related pathways, thanks to the use of language model embeddings to determine spatial layout. This approach ensures that pathways with similar functional roles or shared biological contexts are positioned near each other, facilitating the identification of functionally coherent regions within the map. For instance, in the GBM case study, regions of the Mondrian Map highlighted specific clusters of pathways that were consistently up-regulated or downregulated across different recurrences, offering insights into the tumor’s evolving molecular landscape.

The use of squares to represent fold changes, combined with a color scheme that indicates both the direction of regulation (red for up-regulated, blue for down-regulated) and statistical significance, allows researchers to quickly assess the magnitude and relevance of changes in pathway activity. This visual simplicity contrasts with traditional heatmaps, where the significance and direction of change can be less immediately apparent due to the absence of spatial organization.

Moreover, the colored lines connecting the squares in the Mondrian Map are not merely aesthetic; they convey crucial information about pathway crosstalk. By distinguishing between shortand long-range functional interactions, these lines highlight potential synergistic or antagonistic relationships between pathways that might not be obvious in other forms of visualization. This feature is particularly valuable in understanding complex diseases like GBM, where pathway crosstalk often drives key aspects of tumor progression and resistance to therapy.

The impact of the Mondrian Map extends beyond visualization. Its ability to reveal clusters and interactions within biological networks provides a foundation for more informed bioinformatics analyses, potentially guiding the identification of novel therapeutic targets. By making complex data more accessible and interpretable, the Mondrian Map paves the way for more precise and actionable insights, advancing the field of personalized medicine and network biology.

In summary, the Mondrian Map represents a significant advancement in the visualization of biological networks. It combines the strengths of modern data science techniques with a user-friendly, visually engaging format, making it an invaluable tool for researchers aiming to unravel the complexities of molecular biology and develop more effective therapeutic strategies.

## Acknowledgment

J.C. acknowledges the support of NIH grant awards U54OD036472, U54DK137307, 1OT2OD032742, and R01HL150078.

## Notes

https://github.com/aimed-lab/mondrian-map

### Competing Interest Statement

The authors have declared no competing interest.

### Summary of Updates

The full paper is revised and re-written based on reviewers comments from a top-ranked conference and the workflow is more scientifically rigorous.

