## supplementary-file for "Mondrian Abstraction and Language Model Embeddings for Differential Pathway Analysis"

### Appendix A. Mondrian Abstraction

In Fig. 1, we presented a portrait of Piet Mondrian and one of his iconic art works.

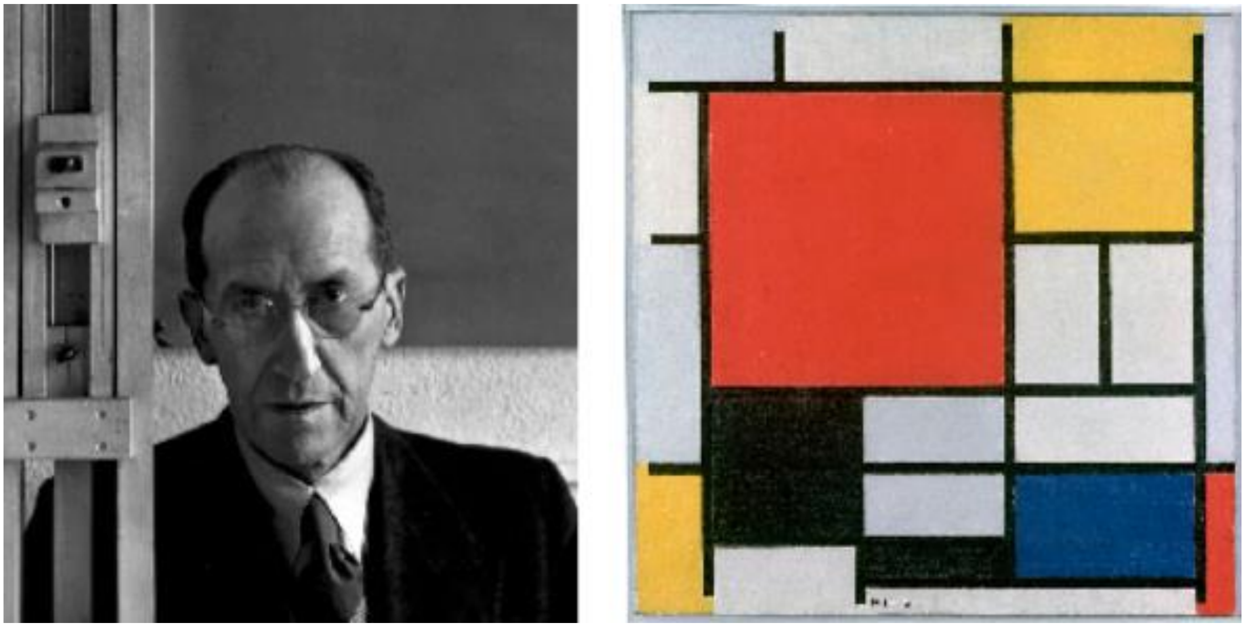

Fig. 1 Mondrian's abstraction. Left: Piet Mondrian (1872–1944), photographed by Arnold Newman in 1942. Right: "Composition with Large Red Plane, Yellow, Black, Grey and Blue" by Mondrian in 1921.

### Appendix B. Prompt Generation

#### Phase I. Prompt for Pathway Description Summarization

**Model:** meta-llama/Meta-Llama-3.1-8B-Instruct.

**Parameters:** max_new_tokens=512, temperature=0.3, do_sample=False.

**Instruction:** Refine the provided biological pathway description by removing all alphanumeric IDs, URLs, and external references. If the description is brief or missing, create a factual summary based on recent literature, avoiding speculation. For lengthy descriptions, produce a concise summary that highlights the key functions and significance of the pathway. Provide only the refined text in one coherent paragraph of up to 300 words, omitting headers and details of deletions.

**Content:** "<Pathway Name>: <Pathway Description>"

**Example:** Sudden infant death syndrome (SIDS) susceptibility pathways: In this model, we provide an integrated view of Sudden Infant Death Syndrome (SIDS) at the level of implicated tissues, signaling networks and genetics. The purpose of this model is to serve as an overview of research in this field and recommend new candidates for more focused or genome wide analyses. SIDS is the sudden and unexpected death of an infant (less than 1 year of age), almost always during deep sleep, where no cause of death can be found by autopsy. Factors that mediate SIDS are likely to be both biological and behavioral, such as sleeping position, environment and stress during a critical phase of infant development (http://www.nichd.nih.gov/health/topics/Sudden_Infant_Death_Syndrome.cfm). While no clear diagnostic markers currently exist, several polymorphisms have been identified which are significantly over-represented in distinct SIDS ethnic population. The large majority of these polymorphisms exist in genes associated with neuronal signaling, cardiac contraction and inflammatory response. These and other lines of evidence suggest that SIDS has a strong autonomic nervous system component (PMID:12350301, PMID: 20124538). One of the neuronal nuclei most strongly implicated in SIDS has been the raphe nucleus of the brain stem. In this nuclei there are ultrastructural, cellular and molecular changes associated with SIDS relative to controls (PMID:19342987, PMID: 20124538). This region of the brain is responsible for the large majority of neuronal serotonin produced and is functionally important in the regulation of normal cardiopulmonary activity, sleep and thermoregulation (see associated references).

Genes associated with serotonin synthesis and receptivity have some of the strongest genetic association with SIDS. Principle among these genes the serotonin biosynthetic enzyme TPH2, the serotonin transporter SLC6A4 and the serotonin receptor HTR1A. SLC6A4 exhibits decreased expression in the raphe nucleus of the medulla oblongata and polymorphisms specifically associated with SIDS (PMID:19342987). In 75% of infants with SIDS, there is decreased HTR1A expression relative to controls along with an increase in the number of raphe serotonin neurons (PMID:19342987). Over-expression of the mouse orthologue of the HTR1A gene in the juvenile mouse medulla produces an analogous phenotype to SIDS with death due to bradycardia and hypothermia (PMID:18599790). These genes as well as those involved in serotonin synthesis are predicted to be transcriptionally regulated by a common factor, FEV (human orthologue of PET-1). PET-1 knock-out results in up to a 90% loss of serotonin neurons

(PMID:12546819), while polymorphisms in FEV are over-represented in African American infants with SIDS. In addition to FEV, other transcription factors implicated in the regulation of these genes (Putative transcriptional regulators (TRs)) and FEV are also listed (see associated references). In addition to serotonin, vasopressin signaling and its regulation by serotonin appear to be important in a common pathway of cardiopulmonary regulation (PMID:2058745). A protein that associates with vasopressin signaling, named pituitary adenylate cyclase-activating polypeptide (ADCYAP1), results in a SIDS like phenotype, characterized by a high increase in spontaneous neonatal death, exacerbated by hypothermia and hypoxia (PMID:14608012), when disrupted in mice. Protein for this gene is widely distributed throughout the central nervous system (CNS), including autonomic control centers (PMID:12389210). ADCYAP1 and HTR1A are both predicted to be transcriptionally regulated by REST promoter binding. Regulation of G-protein coupled signaling pathways is illustrated for these genes, however, it is not clear whether ADCYAP1 acts directly upon raphe serotonin neurons.

Another potentially important class of receptors in SIDS is nicotine. Receptors for nicotine are expressed in serotonin neurons of the raphe throughout development (PMID:18986852). Application of nicotine or cigarette smoke is sufficient to inhibit electrical activity of raphe serotonin neurons (PMID:17515803) and chronic nicotine infusion in rats decreases expression of SLC6A4 (PMID:18778441). Furthermore, nicotine exposure reduces both HTR1A and HTR2A immunoreactivity in several nuclei of the brainstem (PMID:17451658).

In addition to CNS abnormalities, several studies have identified a critical link between cardiac arrhythmia (long QT syndrome) and SIDS (PMID:18928334). A number of genetic association studies identified functionally modifying mutations in critical cardiac channels in as many as 10% of all SIDS cases (PMID:18928334). These mutations have been predicted to predispose infants for long QT syndrome and sudden death. The highest proportion of SIDS associated mutations (both inherited and sporadic) is found in the sodium channel gene SCN5A. Examination of putative transcriptional regulators for these genes, highlights a diverse set of factors as well as a relatively common one (SP1).

Finally, several miscellaneous mutations have been identified in genes associated with inflammatory response and thermoregulation. Infection is considered a significant risk factor for SIDS (PMID:19114412). For inflammatory associated genes, such as TNF alpha, interleukin 10 and complement component 4, many of these mutations are only significant in the presence of infection and SIDS. In addition to these mutations, cerebrospinal fluid levels of IL6 are increased in SIDS cases as well as IL6R levels in the arcuate nucleus of the brain, another major site of serotonin synthesis (PMID:19396608). Genes such as ILR6 and ADCYAP1 are also associated with autoimmune disorders, thus SIDS may also be associated with autoinflammation of autonomic centers in the brain. Regulation of thermogenesis by brown adipose tissue has been proposed be an important component of SIDS, given that SIDS incidence is highest in the winter time and that animal models of SIDS demonstrate variation in body temperature. Interestingly, activation of raphe HTR1A decreases both shivering and peripheral vasoconstriction in piglets (18094064). Although a putative significant polymorphism was identified in the thermoregulator gene HSP60, this only occurred in one SIDS case. It is important to note that in the large majority of all these studies, sleeping position and smoking were among the most significant risk factors for SIDS.

In loving memory of Milo Salomonis (http://www.milosalomonis.org).

Proteins on this pathway have targeted assays available via the [https://assays.cancer.gov/available_assays?wp_id=WP706 CPTAC Assay Portal].

**Generated Summary:** Sudden infant death syndrome (SIDS) is the sudden and unexpected death of an infant under one year of age, often occurring during deep sleep, with no identifiable cause of death. Research suggests that SIDS is influenced by both biological and behavioral factors, including sleeping position, environment, and stress during a critical phase of infant development. Genetic studies have identified several polymorphisms associated with SIDS, primarily in genes related to neuronal signaling, cardiac contraction, and inflammatory response. The autonomic nervous system, particularly the raphe nucleus of the brain stem, is strongly implicated in SIDS, with ultrastructural, cellular, and molecular changes observed in this region. Genes associated with serotonin synthesis and receptivity, such as TPH2, SLC6A4, and HTR1A, have been linked to SIDS, with decreased expression of SLC6A4 and HTR1A observed in SIDS cases. Other factors, including vasopressin signaling, nicotine exposure, and cardiac arrhythmia, have also been implicated in SIDS. In addition, genetic mutations in genes associated with inflammatory response and thermoregulation have been identified, with infection and autoinflammation of autonomic centers in the brain potentially contributing to SIDS. Overall, SIDS is a complex condition influenced by multiple genetic and environmental factors, highlighting the need for further research to understand its underlying mechanisms and develop effective prevention strategies.

**Phase II. Prompt for Embedding Generation**

**Type 1. Gene Symbol**

**Description:** The prompt lists up to 100 gene symbols, sorted by their RP_SCORE in descending order, focusing on the most influential genes while excluding the less influential ones.

**Example:** 'EP300 CREBBP JUN HDAC1 SP1 PRKACA CTNNB1 CREB1 AR VEGFA IL6 TNF NR3C1 NFKB1 IL1B CEBPB HIF1A NGF GAPDH YWHAZ BDNF YWHAE HDAC9 TH PRKACB ESR2 YWHAH CASP3 IL10 EGR1 YWHAQ YWHAG MEF2C SST PPARGC1A HSPD1 PRKAR2A SP3 PRKAR2B RORA ADCYAP1 GRIN1 TPH1 PRKAR1A TCF3 HTR1A YWHAB CREM YBX1 TAC1 THRB IL1A NFKB2 VAMP2 POU5F1 SOX2 CPT1A ASCL1 IL13 MYB SPTBN1 GATA3 HES1 POU2F2 MECP2 PRKAR1B SNTA1 KCNH2 PHOX2A NANOG RET NFYA CHAT RYR2 IL1RN SCN5A CHRNA4 DDC HADHA RUNX3 GJA1 PBX1 MAOA CHRM2 GATA2 KCNQ1 MAP2 PHOX2B IL6R HTR2A ACADM AVP TACR1 SLC6A4 GCK NKX2-2 TP73 HTR3A HADHB MAZ'

**Instruction for llm2vec models:** "Given a list of gene symbols, encode the collective biological significance and pathway associations of these genes:"

**Note:** Along with the main prompt, the instruct models (Mistral-7B and Llama3-8B) require 'instruction' about the task to be performed. This instruction and prompt pair is fed to the model to extract the embeddings.

**Type 2. Gene Description**

**Description:** Similar to Type 1, this includes both Gene Symbols and their Descriptions.

**Example:** 'EP300 (E1A binding protein p300), CREBBP (CREB binding protein), JUN (Jun proto-oncogene, AP-1 transcription factor subunit), HDAC1 (histone deacetylase 1), SP1 (Sp1 transcription factor), PRKACA (protein kinase cAMP-activated catalytic subunit alpha), CTNNB1 (catenin beta 1), CREB1 (cAMP responsive element binding protein 1), AR (androgen receptor), VEGFA (vascular endothelial growth factor A), IL6 (interleukin 6), TNF (tumor necrosis factor), NR3C1 (nuclear receptor subfamily 3 group C member 1), ... ..., GCK (glucokinase), NKX2-2 (NK2 homeobox 2), TP73 (tumor protein p73), HTR3A (5-hydroxytryptamine receptor 3A), HADHB (hydroxyacyl-CoA dehydrogenase trifunctional multienzyme complex subunit beta), MAZ (MYC associated zinc finger protein)'

**Instruction for llm2vec models:** "Given a list of gene symbols and its description, encode the collective biological significance and pathway associations of these genes:"

**Type 3. Pathway Name**

**Description:** The prompt simply consists of the pathway name.

**Example:** Sudden infant death syndrome (SIDS) susceptibility pathways.

**Instruction for llm2vec models:** "Given a pathway name, encode the biological significance and functional associations of this pathway:"

**Type 4. Pathway Description Summary**

**Description:** The prompt consists of the summary generated from *I. Prompt for Pathway Description Summarization*.

**Example:** Sudden infant death syndrome (SIDS) is the sudden and unexpected death of an infant under one year of age, often occurring during deep sleep, with no identifiable cause of death. Research suggests that SIDS is influenced by both biological and behavioral factors, including sleeping position, environment, and stress during a critical phase of infant development. Genetic studies have identified several polymorphisms associated with SIDS, primarily in genes related to neuronal signaling, cardiac contraction, and inflammatory response. The autonomic nervous system, particularly the raphe nucleus of the brain stem, is strongly implicated in SIDS, with ultrastructural, cellular, and molecular changes observed in this region. Genes associated with serotonin synthesis and receptivity, such as TPH2, SLC6A4, and HTR1A, have been linked to SIDS, with decreased expression of SLC6A4 and HTR1A observed in SIDS cases. Other factors, including vasopressin signaling, nicotine exposure, and cardiac arrhythmia, have also been implicated in SIDS. In addition, genetic mutations in genes associated with inflammatory response and thermoregulation have been identified, with infection and autoinflammation of autonomic centers in the brain potentially contributing to SIDS. Overall, SIDS is a complex condition influenced by multiple genetic and environmental factors, highlighting the need for further research to understand its underlying mechanisms and develop effective prevention strategies.

**Instruction for llm2vec models:** "Given a pathway name and its description, encode the biological significance and functional associations of this pathway:"

### Appendix C. Data Preparation for Case Study

**Table 1.** Number of Up and Down-regulated genes per patient profiles from section II (C) (3).

| **Profile** | **# Up-regulated Genes** | **# Down-regulated Genes** |
|---|---|---|
| Baseline R1/TP | 2409 | 362 |
| Baseline R2/TP | 3808 | 315 |
| Aggressive R1/TP | 745 | 923 |
| Aggressive R2/TP | 985 | 1101 |
| Non-aggressive R1/TP | 1727 | 3208 |
| Non-aggressive R2/TP | 2842 | 2583 |

**Table 2.** Number of PAGs and m-type PAG-to-PAG relationships from section II (C) (4).

| Profile | # PAGs | # PAG-to-PAG relationships |
|---|---|---|
| Baseline R1/TP | 302 | 1488 |
| Baseline R2/TP | 533 | 6220 |
| Aggressive R1/TP | 117 | 208 |
| Aggressive R2/TP | 140 | 260 |
| Non-aggressive R1/TP | 460 | 5520 |
| Non-aggressive R2/TP | 560 | 7602 |

Now, we present the resulting data tables from each step, following the data preparation flow diagram.

#### I. Temporal Gene Expression Profile Selection

| | baseline_R1 | baseline_R2 | baseline_TP | aggressive_R1 | aggressive_R2 | aggressive_TP | nonaggressive_R1 | nonaggressive_R2 | nonaggressive_TP |
|---|---|---|---|---|---|---|---|---|---|
| **Gene_symbol** | | | | | | | | | |
| **CMSS1** | 31.207177 | 35.164799 | 24.341754 | 26.446038 | 27.000944 | 32.025694 | 31.395261 | 48.859002 | 44.117700 |
| **STX16** | 35.996421 | 44.218382 | 20.716331 | 99.860372 | 104.422161 | 126.156318 | 20.050733 | 21.480300 | 45.597264 |
| **TLE4** | 20.813337 | 22.299578 | 10.835135 | 23.209358 | 19.561900 | 31.615145 | 72.895632 | 96.452925 | 65.074070 |
| **LRRN1** | 31.667442 | 16.577943 | 53.999371 | 2.167500 | 0.433401 | 5.336610 | 121.089577 | 101.425790 | 137.481300 |
| **YKT6** | 92.464768 | 82.431033 | 75.375286 | 52.902599 | 65.192139 | 59.786585 | 11.058250 | 13.540128 | 9.911100 |
| **...** | ... | ... | ... | ... | ... | ... | ... | ... | ... |
| **UQCRFS1** | 82.462700 | 62.024200 | 65.626250 | 54.498700 | 58.399200 | 50.797400 | 68.277000 | 105.298000 | 60.563400 |
| **C4orf32** | 3.717895 | 4.994350 | 2.917710 | 0.268489 | 0.327187 | 0.151905 | 4.348520 | 4.666130 | 4.126970 |
| **MTRNR2L9** | 36.739885 | 82.480730 | 47.055200 | 0.430776 | 0.616574 | 0.785486 | 111.565000 | 63.872200 | 98.041800 |
| **BMP2** | 5.453945 | 6.221065 | 27.504200 | 0.685032 | 0.462191 | 0.778202 | 36.265900 | 70.232900 | 40.471300 |
| **EP300** | 12.412165 | 13.973550 | 12.963900 | 43.230000 | 41.312600 | 56.060900 | 48.727000 | 37.960800 | 42.483100 |

11090 rows × 9 columns

#### II. Differentially Expressed Gene Selection

| | baseline_R1/TP | baseline_R2/TP |
|---|---|---|
| **Gene_symbol** | | |
| **CMSS1** | 1.282043 | 1.444629 |
| **STX16** | 1.737587 | 2.134470 |
| **TLE4** | 1.920912 | 2.058080 |
| **LRRN1** | 0.586441 | 0.307003 |
| **YKT6** | 1.226725 | 1.093608 |
| **...** | ... | ... |
| **UQCRFS1** | 1.256551 | 0.945113 |
| **C4orf32** | 1.274251 | 1.711736 |
| **MTRNR2L9** | 0.780783 | 1.752850 |
| **BMP2** | 0.198295 | 0.226186 |
| **EP300** | 0.957441 | 1.077882 |

11090 rows × 2 columns

#### III. Gene-set, Network, and Pathway Analysis (GNPA)

| | GS_ID | NAME | SOURCE | GS_SIZE | ORGANISM | DESCRIPTION | LINK | TYPE | MULTI_N | OLAP | COCO_V2 | SIMILARITY_SCORE | pvalue | Rank | pFDR |
|---|---|---|---|---|---|---|---|---|---|---|---|---|---|---|---|
| 0 | WAG002718 | Neutrophil degranulation | WikiPathway_2021 | 474 | Homo sapiens | Neutrophil degranulation | https://www.wikipathways.org/index.php/Pathway... | P | 1333 | 120 | 172.7292014 | .08382793213929303009611849361836230894448 | 1.732402e-44 | 1 | 5.422417e-42 |
| 1 | WAG002628 | VEGFA-VEGFR2 signaling pathway | WikiPathway_2021 | 438 | Homo sapiens | VEGFA-VEGFR2 signaling pathway | https://www.wikipathways.org/index.php/Pathway... | P | 1333 | 97 | 980.8100106 | .0712460408468820826612487529758081271878 | 3.931521e-31 | 2 | 1.226635e-28 |
| 2 | WAG003066 | Endothelin pathway | WikiPathway_2021 | 194 | Homo sapiens | Endothelin pathway | https://www.wikipathways.org/index.php/Pathway... | P | 1333 | 56 | 968.3903924 | .0744733638450988550382938067279983973618 | 1.531451e-24 | 3 | 4.762811e-22 |
| 3 | WAG002670 | TYROBP causal network in microglia | WikiPathway_2021 | 60 | Homo sapiens | TYROBP causal network in microglia | https://www.wikipathways.org/index.php/Pathway... | P | 1333 | 32 | 221.1523998 | .1166432839698623018965965185762535723565 | 3.132260e-24 | 4 | 9.710006e-22 |
| 4 | WAG002820 | Ebola virus pathway in host | WikiPathway_2021 | 131 | Homo sapiens | Ebola virus pathway in host | https://www.wikipathways.org/index.php/Pathway... | P | 1333 | 45 | 650.3115529 | .082421802271457829082107614969279463787 | 1.841119e-23 | 5 | 5.689056e-21 |

#### IV. Pathway Networks

| | GS_A_ID | GS_A_SIZE | GS_B_ID | GS_B_SIZE | OLAP | SIMILARITY | PVALUE |
|---|---|---|---|---|---|---|---|
| 0 | WAG003212 | 430 | WAG003224 | 153 | 55 | .15522875816993464 | 149.90937335538615 |
| 1 | WAG002596 | 246 | WAG002683 | 29 | 16 | .15976567700705632 | 62.03314298178884 |
| 2 | WAG002596 | 246 | WAG002118 | 55 | 31 | .20457912457912455 | 120.66116349772959 |
| 3 | WAG002596 | 246 | WAG003300 | 33 | 19 | .17361305361305365 | 74.66521990991453 |
| 4 | WAG002414 | 90 | WAG002096 | 63 | 18 | .16380952380952382 | 73.94134202903723 |

#### V. Differential Pathway Analysis

| | GS_ID | wFC | pFDR |
|---|---|---|---|
| 0 | WAG001976 | 1.3186 | 0.006331 |
| 1 | WAG001977 | 1.6080 | 0.075830 |
| 2 | WAG002000 | 2.0673 | 0.000030 |
| 3 | WAG002011 | 1.0627 | 0.240331 |
| 4 | WAG002018 | 1.7749 | 0.084468 |

#### VI. Pathway Embeddings

Utilizing language models to generate pathway embeddings described in section II A and Appendix C, we get one final data table for each profile with top 10 pathways ready for Mondrian Map Visualization. The table below is for Baseline cohort R1/TP. We have 5 similar tables for 5 other profiles.

| | GS_ID | wFC | pFDR | x | y |
|---|---|---|---|---|---|
| 0 | WAG002133 | 1.6183 | 3.724584e-43 | 607.298815 | 220.621129 |
| 1 | WAG002628 | 2.0947 | 3.882609e-43 | 200.916562 | 272.423214 |
| 2 | WAG002134 | 1.6776 | 1.419164e-41 | 704.595680 | 265.673499 |
| 3 | WAG003066 | 2.5782 | 2.237119e-30 | 185.874616 | 133.851282 |
| 4 | WAG003212 | 2.2273 | 3.933392e-30 | 180.720278 | 392.922252 |
| 5 | WAG002596 | 1.7945 | 2.100707e-29 | 287.372243 | 443.144485 |
| 6 | WAG002718 | 1.6066 | 2.283309e-27 | 775.578285 | 92.995014 |
| 7 | WAG003032 | 2.3010 | 6.165519e-26 | 446.478214 | 887.448126 |
| 8 | WAG002186 | 1.7709 | 5.848443e-23 | 93.562092 | 282.262669 |
| 9 | WAG002879 | 1.5897 | 9.164786e-23 | 70.818373 | 160.476481 |

#### Appendix D. Pathway Embeddings

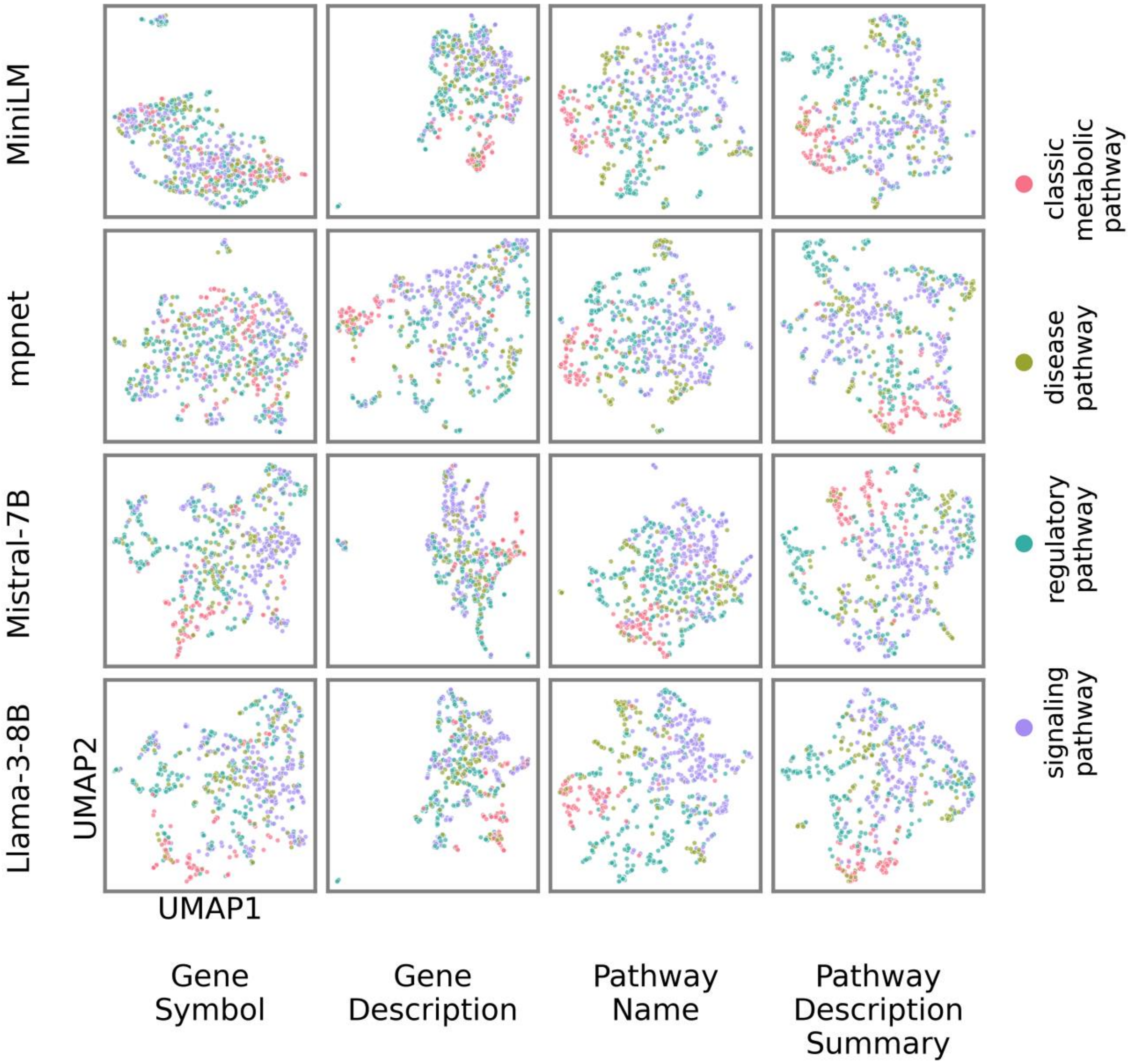


Fig. 2 UMAP projection for embeddings. It is more converged compared to t-SNE projection shown in the paper.

model: Llama-3-8B, type: Pathway Name  (embedding=0.83, umap=0.71, tsne=0.73)

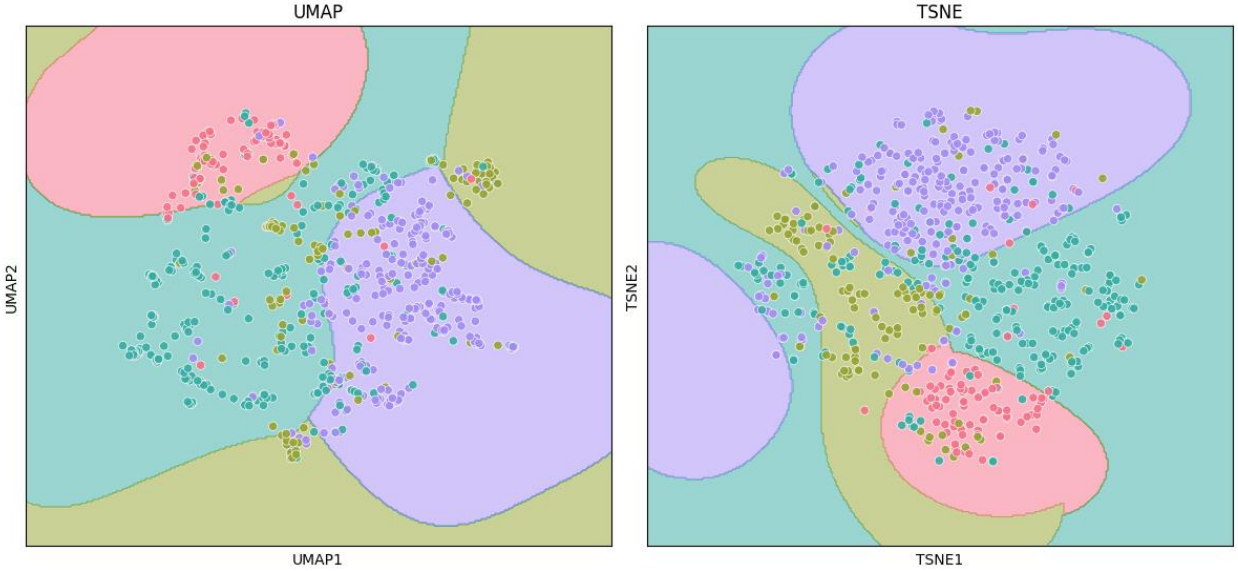


model: Llama-3-8B, type: Pathway Description Summary  (embedding=0.79, umap=0.7, tsne=0.72)

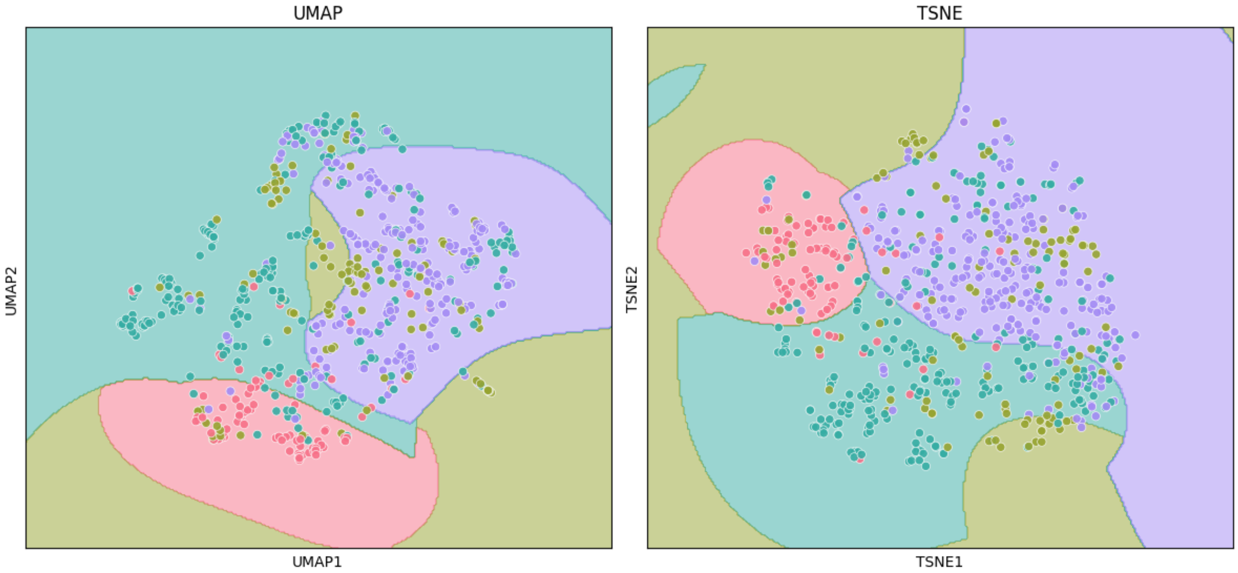

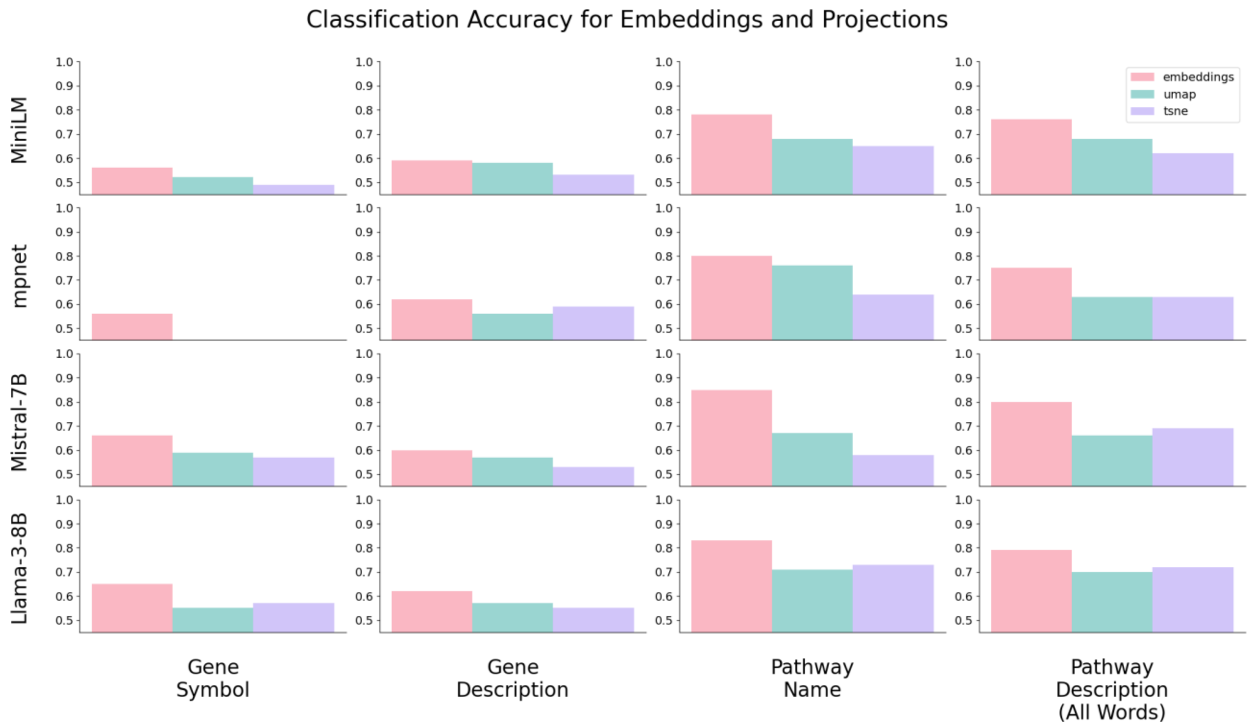

Classification Accuracy for Embeddings and Projections
embeddings
umap
tsne
MiniLM
mpnet
Mistral-7B
Llama-3-8B
1.0
0.9
0.8
0.7
0.6
0.5
Gene Symbol
Gene Description
Pathway Name
Pathway Description (All Words)

### Appendix E. Pathway IDs, Names and Descriptions Visualized in Mondrian Maps

Out[26]:

| | ID | Pathway Name | Pathway Description |
|---|---|---|---|
| **0** | 2048 | Generic transcription pathway | The transcriptional regulation of gene expression in eukaryotic cells is a complex process involving multiple components and mechanisms. The general principles of transcription regulation have been revealed through detailed studies of various eukaryotic systems. The core promoter is a crucial region where the RNA polymerase II (Pol II) and general factors bind to specific DNA motifs, including the TATA box, Initiator motif, and Downstream Promoter Element. The proximal promoter region, located immediately upstream of the core promoter, contains functional DNA binding sites for transcription activator and repressor proteins, which are often cell- or tissue-specific in expression. Distal enhancer regions, located far from the promoter, also play a crucial role in regulating gene expression, often in a cell- or tissue-specific pattern. The specific combination of transcription activator and repressor binding sites within the proximal promoter and/or distal enhancer regions provides a "combinatorial transcription code" that mediates cell- or tissue-specific expression of the associated target gene. The assembly of co-activator and co-repressor complexes on DNA-bound transcription factors is essential for regulating target gene transcription, with co-activators typically containing histone acetyltransferase activity and co-repressors containing histone deacetylase activity. The Mediator complex, a large co-activator complex, bridges between the basal transcription factors and tissue-specific transcription factors, while the TBP-associated factors (TAFs) assemble on TBP, bound to the TATA box. Specific co-activator complexes for DNA-binding transcription factors, such as DRIP, TRAP, and ARC, have been identified, and these complexes are essential for regulating target gene transcription. The Notch signaling pathway is a well-studied example of cell-specific regulation of gene transcription, where the CSL protein is a bifunctional DNA-binding transcription factor that mediates repression in the absence of Notch signaling and activation in the presence of Notch signaling. |
| **1** | 2068 | Mitotic G2-G2/M phases | The mitotic G2 phase is the second growth phase in the eukaryotic mitotic cell cycle, occurring between the completion of DNA synthesis and the beginning of mitosis. During this phase, the cell's cytoplasmic content increases in preparation for cell division. As the cell transitions from G2 to M, duplicated centrosomes mature and separate, and CDK1:cyclin B complexes become active. This activation sets the stage for spindle assembly and chromosome condensation, which are critical events in the prophase of mitosis. These processes are essential for the accurate segregation of chromosomes during cell division, ensuring the integrity of the genetic material in the daughter cells. The proper progression through the G2 phase and the subsequent transition to mitosis are tightly regulated by a complex interplay of molecular mechanisms, ultimately leading to the successful completion of cell division. |
| **2** | 2133 | miR-targeted genes in lymphocytes | The pathway involves microRNAs (miRs) that target specific genes in lymphocytes, a type of immune cell. Lymphocytes play a crucial role in the adaptive immune response, and miRs are small non-coding RNAs that regulate gene expression by binding to messenger RNA (mRNA) and preventing its translation. In lymphocytes, miRs are involved in various processes, including cell proliferation, differentiation, and survival. They regulate the expression of genes involved in immune responses, such as cytokine production and antigen presentation. The pathway highlights the complex interactions between miRs and their target genes in lymphocytes, which are essential for maintaining immune homeostasis and responding to pathogens. By regulating gene expression, miRs help to fine-tune the immune response, preventing excessive inflammation and tissue damage. This pathway provides a comprehensive overview of the miR-targeted genes in lymphocytes, highlighting their significance in immune function and disease. |
| **3** | 2134 | miR-targeted genes in muscle cell | The muscle cell pathway involves the regulation of gene expression by microRNAs (miRs), which are small non-coding RNAs that play a crucial role in post-transcriptional gene regulation. In muscle cells, specific miRs target and regulate the expression of genes involved in various cellular processes, including muscle development, differentiation, and function. These miR-targeted genes are essential for maintaining muscle homeostasis and responding to physiological demands. The regulation of these genes by miRs is a complex process that involves the binding of miRs to the 3' untranslated region (UTR) of target mRNAs, leading to mRNA degradation or translational repression. This pathway highlights the importance of miR-mediated regulation in muscle cells, and its dysregulation has been implicated in various muscle-related disorders, such as muscular dystrophy and muscle atrophy. Understanding the muscle cell pathway can provide insights into the molecular mechanisms underlying muscle development and disease, and may lead to the identification of novel therapeutic targets for muscle-related disorders. |

| | ID | Pathway Name | Pathway Description |
|---|---|---|---|
| **4** | 2173 | Allograft Rejection | The allograft rejection pathway is a fundamental adaptive immune response that leads to the destruction of transplanted tissues. Antigen-presenting cells, either from the donor or recipient, activate naive T cells, resulting in the maturation of CD8+ and CD4+ T cells. CD8+ T cells induce apoptosis in donor cells, while CD4+ T cells differentiate into various subsets, including TH1, TH2, T17, and Treg cells. Activated TH1 cells produce pro-inflammatory cytokines, such as TNFA and NO, which damage donor graft cells through cytotoxicity. In contrast, TH2 cells activate B cells, leading to the production of IgG antibodies and the complement cascade, which contributes to acute and chronic antibody-mediated rejection. The complement cascade is a key therapeutic target, with inhibitors like Eculizumab and YCF preventing its activation. Additionally, other immune suppressants, such as corticosteroids, Belatacept, and CTLA4, inhibit T-cell activation and pro-inflammatory cytokines, contributing to immune suppression and preventing allograft rejection. |
| **5** | 2186 | Brain-derived neurotrophic factor (BDNF) signaling pathway | Brain-derived neurotrophic factor (BDNF) plays a crucial role in the growth, differentiation, plasticity, and survival of neurons, as well as in various physiological processes such as energy metabolism, behavior, mental health, learning, memory, stress, pain, and apoptosis. BDNF is implicated in various neuronal disorders, including Alzheimer's disease, Huntington's disease, depression, and bipolar disorder. BDNF binds to the tropomyosin-related kinase B (TrkB) receptor, triggering a signaling cascade that involves the activation of multiple downstream targets, including Shp2, Shc, and PLC-gamma. This leads to the activation of various signaling pathways, including PLC/PKC, PI3K/Akt, Ras/Erk, AMPK/ACC, and NFκB, which regulate processes such as synaptic plasticity, cell survival, and protein synthesis. BDNF also interacts with the p75 neurotrophin receptor (p75NTR), which can lead to apoptosis through the activation of JNK. The PI3K/Akt pathway, activated by BDNF/TrkB interaction, inhibits cell apoptosis and promotes neuronal survival. BDNF signaling also regulates the surface expression of AMPA and NMDA receptors, and the expression of genes involved in processes such as dendrite differentiation and calcification of cementoblast-like cells. Overall, BDNF plays a critical role in maintaining neuronal health and function, and its dysregulation is implicated in various neurological disorders. |
| **6** | 2190 | Spinal cord injury | Spinal cord injury is a complex process involving the regulation of gene expression and signaling in various cell types, including motor neurons, oligodendrocytes, microglia, and astrocytes. This leads to immediate immune responses that last several weeks, characterized by inflammation, excitotoxicity, cell death, and the formation of a glial scar, ultimately suppressing axonal regeneration. The injury triggers the release of chemoattractants and cytokines, which activate neutrophils and recruit immune cells, including B and T cells, and macrophages. This immune response results in the proliferation of astrocytes and microglia, leading to apoptosis and necrosis of oligodendrocytes and neurons. The expression of cell cycle genes is increased, contributing to the proliferation of these cells. An example therapy for spinal cord injury is the administration of immunosuppressants, such as FK506, which offers neuroprotection. The goal of such therapies is to mitigate the immune response and promote a conducive environment for axonal regeneration and tissue repair. |
| **7** | 2380 | Nuclear receptors meta-pathway | Nuclear receptors are a class of proteins found within cells that sense steroid and thyroid hormones, as well as certain other molecules, to regulate gene expression. These receptors work with other proteins to control the development, homeostasis, and metabolism of the organism by directly binding to DNA and regulating the expression of adjacent genes. This process, known as transcriptional regulation, is typically initiated by the binding of a ligand to the nuclear receptor, resulting in a conformational change that activates the receptor and leads to the up- or down-regulation of gene expression. As transcription factors, nuclear receptors play crucial roles in both embryonic development and adult homeostasis, making them essential for maintaining the balance of various physiological processes. Their unique ability to directly interact with and control genomic DNA sets them apart from other classes of receptors, underscoring their significance in cellular regulation. |
| **8** | 2393 | Focal adhesion | Focal adhesions are complex structures formed at the contact points between cells and the extracellular matrix, playing crucial roles in various biological processes such as cell motility, proliferation, differentiation, gene expression regulation, and cell survival. These structures consist of bundles of actin filaments anchored to transmembrane integrin receptors through a multi-molecular complex of junctional plaque proteins. Focal adhesions contain both structural components, such as proteins linking membrane receptors to the actin cytoskeleton, and signaling molecules, including protein kinases, phosphatases, and adapter proteins. The non-receptor tyrosine kinase activities of FAK and src proteins, along with the adaptor protein functions of FAK, src, and Shc, initiate downstream signaling events that lead to the reorganization of the actin cytoskeleton, changes in cell shape and motility, and modulation of gene expression. This signaling pathway exhibits significant crosstalk with growth factor-mediated signaling, highlighting the importance of focal adhesions in regulating cellular behavior and function. |

| | ID | Pathway Name | Pathway Description |
|---|---|---|---|
| **9** | 2483 | Assembly of the primary cilium | The primary cilium is a non-motile sensory organelle present in a single copy at the apical surface of most quiescent cells, playing crucial roles in signaling and development. Cilium biogenesis involves the anchoring of the basal body near the plasma membrane and the subsequent polymerization of the microtubule-based axoneme and extension of the plasma membrane. The ciliary membrane is distinct from the bulk cytoplasm and plasma membrane, established and maintained during biogenesis by the formation of a ciliary transition zone. Ciliary components are targeted to the ciliary base and transported to the ciliary tip by intraflagellar transport (IFT), a motor-driven process requiring kinesin-2 and dynein-2 motors and the IFT complex. The primary cilium undergoes continuous turnover of tubulin at the tip, requiring the IFT machinery for maintenance as well as biogenesis. The importance of the primary cilium is highlighted by the wide range of defects and disorders, collectively known as ciliopathies, that arise from mutations in genes encoding components of the ciliary machinery. |
| **10** | 2596 | MAPK signaling pathway | The mitogen-activated protein kinase (MAPK) cascade is a highly conserved module that plays a crucial role in various cellular functions, including cell proliferation, differentiation, and migration. In mammals, there are four distinct groups of MAPKs: ERK-1/2, JNK1/2/3, p38 proteins, and ERK5, which are activated by specific MAPKKs. Each MAPKK can be activated by multiple MAPKKKs, increasing the complexity and diversity of MAPK signaling. This allows cells to respond to various stimuli, such as growth factors and pro-inflammatory signals, through distinct MAPKKKs. The MAPK cascade is essential for regulating cellular processes, and its dysregulation has been implicated in various diseases, including cancer. |
| **11** | 2628 | VEGFA-VEGFR2 signaling pathway | The VEGFA-VEGFR2 signaling pathway plays a central role in angiogenesis, the formation of new blood vessels from pre-existing vasculature, which is crucial for various physiological conditions such as embryogenesis, wound healing, and is a hallmark of pathological conditions like tumorigenesis. Vascular endothelial growth factor (VEGF) is the primary angiogenic growth factor that modulates neovascularization. VEGF binds to VEGFR2 on the surface of endothelial cells, leading to dimerization and auto-phosphorylation of specific tyrosine residues, which activates multiple downstream signaling cascades. This activation promotes endothelial cell proliferation, migration, and tube formation, essential for angiogenesis. The PI3K-AKT-mTOR signaling pathway, regulated by VEGFR2, is involved in cell survival, proliferation, and anti-apoptotic functions, while PLCÎ³-mediated activation of PKC and downstream induction of ERK and other PKC-dependent pathways contribute to cell proliferation. Additionally, VEGFA/VEGFR2 signaling induces endothelial cell migration through activation of p38MAPK and FAK, which is crucial for directed migration. The VEGFA/VEGFR2 signaling network is a complex interplay of protein-protein interactions, enzyme-catalyzed events, and gene regulation, which modulates angiogenesis and is essential for various physiological and pathological processes. |
| **12** | 2659 | Focal adhesion: PI3K-Akt-mTOR-signaling pathway | Focal adhesions play a crucial role in various biological processes, including cell motility, proliferation, differentiation, gene expression regulation, and cell survival. These structures, formed at the cell-extracellular matrix contact points, consist of bundles of actin filaments anchored to trans-membrane receptors of the integrin family through a complex of multiple proteins. Focal adhesions not only provide a structural link between membrane receptors and the actin cytoskeleton but also contain signaling molecules that initiate downstream events. The non-receptor tyrosine kinase activities of FAK and src proteins, along with the adaptor protein functions of FAK, src, and Shc, are essential for initiating these signaling events. The resulting reorganization of the actin cytoskeleton is vital for changes in cell shape and motility, as well as gene expression. This pathway is significant in understanding the mechanisms underlying cell behavior and its implications in various diseases, including cancer. |
| **13** | 2670 | TYROBP causal network in microglia | The TYROBP causal network in microglia plays a crucial role in the regulation of immune responses in the brain. Microglia, the primary immune cells of the central nervous system, express TYROBP, also known as DAP12, which is an adaptor protein involved in signaling through various receptors. The TYROBP network is essential for the proper functioning of microglia, influencing their activation, proliferation, and survival. This pathway is also involved in the regulation of inflammatory responses, with TYROBP signaling contributing to the production of pro-inflammatory cytokines and the activation of immune cells. The TYROBP network has been implicated in various neurological disorders, including Alzheimer's disease, Parkinson's disease, and multiple sclerosis, where microglial dysfunction is thought to play a key role. Further research is needed to fully understand the complex interactions within the TYROBP network and its significance in maintaining immune homeostasis in the brain. |

| | ID | Pathway Name | Pathway Description |
|---|---|---|---|
| **14** | 2718 | Neutrophil degranulation | Neutrophils, the most abundant leukocytes, play a crucial role in defending the body against invading microorganisms. In response to infection, they migrate towards the inflammatory focus, mobilizing granules that fuse with the cell membrane or phagosomal membrane, resulting in the exocytosis or exposure of membrane proteins. Neutrophil granules contain antimicrobial and proteolytic substances, as well as membrane proteins that change how the neutrophil responds to its environment. Primed neutrophils actively secrete cytokines and other inflammatory mediators, and can present antigens via MHC II, stimulating T-cells. Granules form during neutrophil differentiation, with subtypes distinguished by their content, but sharing similarities in structure and composition. The classical granule subsets include Azurophil, secondary, and gelatinase granules, as well as storage vesicles formed by endocytosis, which contain cell-surface markers and extracellular proteins. These granules are highly exocytosable, allowing neutrophils to rapidly respond to and eliminate pathogens. |
| **15** | 2783 | Cilium Assembly | Cilia are membrane-covered organelles extending from the surface of eukaryotic cells, distinguished by the structure of their microtubule-based axonemes. Four main types of cilia have been identified in humans, including motile and non-motile cilia with distinct axoneme structures. The primary cilium, a non-motile sensory organelle, is required for the transduction of external signals, including growth factors, hormones, and morphogens, and is essential for signaling pathways mediated by various receptors. Cilium biogenesis involves the anchoring of the basal body near the plasma membrane, polymerization of the axoneme, and extension of the plasma membrane. The ciliary membrane is distinct from the bulk cytoplasm and plasma membrane, established and maintained during biogenesis by the formation of a ciliary transition zone and intraflagellar transport (IFT). IFT is a motor-driven process that targets ciliary components from the secretory system to the ciliary tip, requiring kinesin-2 and dynein-2 motors and the IFT complex. The primary cilium undergoes continuous turnover of tubulin at the tip, requiring the IFT machinery for maintenance as well as biogenesis. Defects in ciliary components can lead to ciliopathies, a wide range of disorders resulting from mutations in genes encoding ciliary machinery components. |
| **16** | 2805 | PI3K-Akt signaling pathway | The PI3K-Akt signaling pathway is a crucial cellular mechanism that regulates various fundamental processes, including apoptosis, protein synthesis, metabolism, and cell cycle progression. It is activated by diverse cellular stimuli and toxic insults, leading to the phosphorylation and activation of Akt by PI3K. Once active, Akt exerts its effects by phosphorylating a range of substrates, thereby controlling downstream cellular processes. This pathway plays a significant role in maintaining cellular homeostasis and is often dysregulated in various diseases, including cancer. The PI3K-Akt pathway is also involved in insulin signaling, glucose metabolism, and cell survival, highlighting its importance in maintaining proper cellular function. |
| **17** | 2820 | Ebola virus pathway in host | The Ebola virus infection pathway in humans involves the attachment of the virus to the plasma membrane of host cells, primarily macrophages and dendritic cells, although it can infect a wide range of cells. The viral glycoprotein induces endocytosis, a process facilitated by membrane proteins, allowing the virus to penetrate the cell. Within the endosomal compartments, viral glycoproteins are cleaved and fused to the endosomal membrane, resulting in the uncoating of viral particles into the cytoplasm. Once inside the cell, the virus begins to replicate and suppresses the host's immune response. The newly created viruses are released from host cells through various mechanisms, including cell lysis, waiting for cell death, or budding off through the cell membrane. This complex process ultimately leads to the spread of the virus within the host, highlighting the need for a comprehensive understanding of the Ebola virus infection pathway to develop effective treatments and prevention strategies. |
| **18** | 2876 | Ciliary landscape | The ciliary landscape pathway is a complex network of proteins and interactions that play a crucial role in ciliary function. Cilia are essential cellular structures involved in sensing the environment, regulating cell signaling, and maintaining proper cellular function. The pathway is composed of 1319 proteins and 4905 interactions, with many proteins having multiple roles and connections. Dysfunction in ciliary function is a hallmark of ciliopathies, a group of diseases that affect various organs and tissues, including the kidneys, eyes, and brain. Ciliopathies include polycystic kidney disease, Usher syndrome, Bardet-Biedl syndrome, Meckel-Gruber syndrome, and Jeune syndrome, among others. These diseases are often characterized by impaired ciliary motility, structure, or signaling, leading to a range of symptoms and complications. Understanding the ciliary landscape pathway is essential for developing new therapeutic strategies to treat ciliopathies and other related disorders. |

| | ID | Pathway Name | Pathway Description |
|---|---|---|---|
| **19** | 2879 | EGF/EGFR signaling pathway | The EGF/EGFR signaling pathway plays a crucial role in regulating cell growth, differentiation, migration, adhesion, and survival. Upon binding of epidermal growth factor (EGF) to the epidermal growth factor receptor (EGFR), the receptor undergoes dimerization, activation of intrinsic kinase activity, and subsequent autophosphorylation. This triggers the recruitment of various cytoplasmic proteins, including adapter proteins, enzymes, and substrates, which transduce and regulate the EGFR function. The pathway activates multiple signaling modules, including the RAS, PLC, STAT, FAK, JNK, p38MAPK, and ERK5 modules, leading to the activation of transcriptional regulators and the induction of cell growth and proliferation. The pathway also regulates cell survival through the activation of AKT and the PI3K/AKT pathway. EGFR signaling is tightly regulated by various proteins, including CBL, CSK, PKC, and PTEN, which promote endocytosis or reduction in EGFR activity or its signaling mediators. Aberrant EGFR signaling has been implicated in oncogenesis, making it a promising target for cancer therapy. |
| **20** | 2896 | G alpha (i) signaling events | G alpha (i) signaling plays a crucial role in inhibiting the cAMP-dependent pathway by suppressing adenylate cyclase activity, thereby reducing cAMP production and subsequent activation of cAMP-dependent protein kinases. This inhibition has significant downstream effects on cellular processes. In addition to its role in cAMP regulation, G alpha (i) also activates the protein tyrosine kinase c-Src, which is involved in various cellular signaling pathways. The activity of G alpha (i) can be modulated by Regulator of G-protein Signalling (RGS) proteins, which regulate the G protein's activity and influence its downstream effects. The G alpha (i) signaling pathway is essential for maintaining cellular homeostasis and responding to various extracellular stimuli, and its dysregulation has been implicated in various diseases, including cancer and cardiovascular disorders. |
| **21** | 2973 | Fragile X syndrome | Fragile X syndrome is a genetic disorder caused by a mutation in the FMR1 gene, leading to the most common form of inherited intellectual disability and autism spectrum disorder. Characteristic physical features include macro-orchidism in males, a long and narrow face, large and protruding ears, and hyperextensible joints. Common comorbidities include neuropsychiatric disorders such as hyperactivity, depression, and anxiety. The mutation disrupts production of the fragile mental retardation protein (FMRP), which regulates de novo protein synthesis and synaptic plasticity by acting as a translational repressor for target mRNAs. FMRP, along with the mTOR and ERK pathways, influences the expression of target mRNAs through stimulation of Group I metabotropic glutamate receptors, ultimately affecting the internalization of α-amino-3-hydroxy-5-methyl-4-isoxazolepropionic acid receptors and long-term depression. The lack of FMRP leads to exaggerated long-term depression, a form of synaptic plasticity involved in learning and memory, which is a primary contributor to the pathogenesis of fragile X syndrome. |
| **22** | 3031 | IL-18 signaling pathway | Interleukin-18 (IL-18) is a cytokine that plays a significant role in the activation of various hematopoietic cell types, mediating both Th1 and Th2 responses. It is the primary inducer of interferon-γ in these cells and is involved in the production and release of several cytokines, chemokines, and cellular adhesion molecules. IL-18's biological activity is mediated through its binding to the IL-18 receptor complex, leading to the activation of nuclear factor-kB (NF-kB) and the subsequent production of proinflammatory cytokines. Additionally, IL-18 activates mitogen-activated protein kinases (MAPKs) and PI3K/AKT signaling modules in certain cell types, contributing to its role in host defense, inflammation, and tissue regeneration. As a vital component of the immunomodulatory cytokine networks, IL-18 signaling plays a crucial part in the body's response to pathogens and injury, highlighting its importance in maintaining immune homeostasis. |
| **23** | 3032 | PodNet: protein-protein interactions in the podocyte | The podocyte is a specialized cell type in the kidney that plays a crucial role in maintaining the integrity of the glomerular filtration barrier. The podocyte's function is supported by a complex network of protein-protein interactions, which are essential for its structure and function. These interactions involve various proteins that are localized to the podocyte's plasma membrane, cytoskeleton, and nucleus. The podocyte's protein-protein interaction network, also known as PodNet, is a critical component of its cellular architecture and is involved in regulating its morphology, adhesion, and signaling properties. Abnormalities in PodNet have been implicated in various kidney diseases, including focal segmental glomerulosclerosis and minimal change disease. Further research on PodNet is necessary to understand its mechanisms and to identify potential therapeutic targets for treating podocyte-related disorders. |

| | ID | Pathway Name | Pathway Description |
|---|---|---|---|
| **24** | 3063 | AXL signaling pathway | The Axl signaling pathway is a critical biological process mediated by the Axl receptor, a member of the TAM family of receptor tyrosine kinases. Axl is a transmembrane protein with an extracellular domain containing immunoglobulin-like and fibronectin type-III domains, a single-pass transmembrane domain, and an intracellular protein tyrosine kinase domain. The receptor is activated by its ligand, growth arrest-specific protein 6 (Gas6), and can also be activated independently in response to oxidative stress and overexpression. Axl is extensively expressed in various tissues and plays a crucial role in regulating physiological processes such as cell proliferation, survival, motility, aggregation, and anti-inflammation. The aberrant expression and prolonged activation of Axl have been linked to various disease conditions, including cancer, chronic immune disorders, and cardiovascular diseases. Axl activation contributes to various downstream signaling pathways, including PI3K/AKT, MAPK/ERK, JAK/STAT, and PI3K/AKT/mTOR, which regulate cell proliferation, migration, and survival. The Axl signaling pathway has significant biomedical importance, and its dysregulation has been implicated in the development and progression of various diseases, including colorectal cancer. |
| **25** | 3066 | Endothelin pathway | The endothelin pathway is a complex network of signaling events mediated by the endothelin peptide, a potent vasoconstrictor. Endothelin plays a crucial role in regulating various physiological and pathophysiological processes in mammals, exhibiting both beneficial and detrimental effects. The pathway is involved in vasoconstriction, which can impact blood pressure and cardiovascular function. Endothelin also regulates gene expression, protein expression, and molecular associations, influencing cellular processes such as cell growth, differentiation, and survival. Additionally, endothelin is implicated in various disease states, including hypertension, cardiovascular disease, and cancer. The pathway's intricate mechanisms involve multiple activation and inhibition events, post-translational modifications, and protein interactions, highlighting its significance in maintaining cellular homeostasis and its potential as a therapeutic target for various diseases. |
| **26** | 3196 | Burn wound healing | Burn wound healing is a complex process involving multiple cellular and molecular mechanisms. The initial inflammatory response is triggered by the release of cytokines and chemokines, such as IL-1β, TNF-α, and IL-6, which recruit immune cells to the wound site. This is followed by the proliferation phase, where fibroblasts and keratinocytes migrate to the wound area, producing extracellular matrix and promoting tissue regeneration. Growth factors like PDGF, TGF-β, and VEGF play crucial roles in regulating cell proliferation, differentiation, and angiogenesis. The final remodeling phase involves the deposition of collagen and other matrix components, leading to the restoration of tissue strength and function. The healing process is also influenced by various signaling pathways, including the MAPK and PI3K/AKT pathways, which regulate cell survival, migration, and differentiation. Additionally, the wound microenvironment, including the presence of bacteria and other pathogens, can significantly impact the healing process. Overall, burn wound healing is a highly coordinated and dynamic process, involving the interplay of multiple cellular and molecular mechanisms to restore tissue integrity and function. |
| **27** | 3212 | Malignant pleural mesothelioma | Malignant pleural mesothelioma is a rare and aggressive form of cancer that affects the pleura, the thin layer of tissue surrounding the lungs. It is primarily caused by exposure to asbestos, a group of naturally occurring minerals that were widely used in construction, insulation, and other industries until their health risks became well-known. The disease typically develops after a long latency period, often 20-50 years, following initial asbestos exposure. Asbestos fibers can cause genetic mutations in the cells lining the pleura, leading to uncontrolled cell growth and tumor formation. The most common symptoms of malignant pleural mesothelioma include shortness of breath, chest pain, and fatigue, although some patients may not experience any noticeable symptoms until the disease is advanced. Diagnosis is often made through imaging tests, such as CT scans or X-rays, and biopsy. Treatment options are limited and typically involve a combination of surgery, chemotherapy, and radiation therapy, although the prognosis for patients with malignant pleural mesothelioma remains poor. Research into the disease is ongoing, with a focus on developing more effective treatments and improving patient outcomes. |
| **28** | 3238 | Network map of SARS-CoV-2 signaling pathway | The SARS-CoV-2 signaling pathway is a complex network of protein-protein interactions and downstream molecular events regulated by the virus. This pathway involves various molecular processes, including molecular association, catalysis, and gene regulation, which are crucial for the virus's replication and survival within the host cell. The pathway is characterized by a series of interactions between viral and host proteins, leading to the activation of various signaling cascades that ultimately result in the modulation of host cell functions. Post-translational modifications, such as phosphorylation and ubiquitination, play a significant role in regulating the activity of key proteins involved in this pathway. The SARS-CoV-2 signaling pathway is essential for the virus's ability to manipulate host cell processes, including immune evasion, cell cycle progression, and apoptosis, ultimately contributing to the development of severe respiratory disease. Understanding this pathway is critical for the development of effective therapeutic strategies to combat SARS-CoV-2 infection. |

| | ID | Pathway Name | Pathway Description |
|---|---|---|---|
| **29** | 3247 | Alzheimer's disease | Alzheimer's disease is a complex neurodegenerative disorder characterized by progressive cognitive decline and memory loss. The disease is primarily associated with the accumulation of amyloid-beta plaques and tau protein tangles in the brain, leading to neuronal damage and death. The amyloid precursor protein (APP) is cleaved by enzymes such as beta-secretase and gamma-secretase, resulting in the production of amyloid-beta peptides. These peptides then aggregate and form insoluble fibrils that deposit in the brain, contributing to the development of Alzheimer's disease. The tau protein, on the other hand, is a microtubule-associated protein that becomes hyperphosphorylated and forms neurofibrillary tangles, which are another hallmark of the disease. The exact mechanisms underlying Alzheimer's disease are not fully understood, but it is believed that a combination of genetic, environmental, and lifestyle factors contribute to its development. Research has identified several key pathways and molecules involved in the disease, including the insulin signaling pathway, the Wnt signaling pathway, and the immune response. Understanding these pathways and their interactions may provide insights into the development of effective therapeutic strategies for Alzheimer's disease. |
| **30** | 3294 | Sudden infant death syndrome (SIDS) susceptibility pathways | Sudden infant death syndrome (SIDS) is the sudden and unexpected death of an infant under one year of age, often occurring during deep sleep, with no identifiable cause of death. Research suggests that SIDS is influenced by both biological and behavioral factors, including sleeping position, environment, and stress during a critical phase of infant development. Genetic studies have identified several polymorphisms associated with SIDS, primarily in genes related to neuronal signaling, cardiac contraction, and inflammatory response. The autonomic nervous system, particularly the raphe nucleus of the brain stem, is strongly implicated in SIDS, with ultrastructural, cellular, and molecular changes observed in this region. Genes associated with serotonin synthesis and receptivity, such as TPH2, SLC6A4, and HTR1A, have been linked to SIDS, with decreased expression of SLC6A4 and HTR1A observed in SIDS cases. Other factors, including vasopressin signaling, nicotine exposure, and cardiac arrhythmia, have also been implicated in SIDS. In addition, genetic mutations in genes associated with inflammatory response and thermoregulation have been identified, with infection and autoinflammation of autonomic centers in the brain potentially contributing to SIDS. Overall, SIDS is a complex condition influenced by multiple genetic and environmental factors, highlighting the need for further research to understand its underlying mechanisms and develop effective prevention strategies. |
